# Affective Bias as a Rational Response to the Statistics of Rewards and Punishments

**DOI:** 10.1101/114991

**Authors:** Erdem Pulcu, Michael Browning

## Abstract

Affective bias, the tendency to prioritise the processing of negative relative to positive events, is causally linked to clinical depression. However, why such biases develop or how they may best be ameliorated is not known. Using a computational framework, we investigated whether affective biases may reflect an individual’s estimates of the information content of negative and positive events. During a reinforcement learning task, the information content of positive and negative outcomes was manipulated independently by varying the volatility of their occurrence. Human participants altered the learning rates used for the outcomes selectively, preferentially learning from the most informative. This behaviour was associated with activity of the central norepinephrine system, estimated using pupilometry, for loss outcomes. Humans maintain independent estimates of the information content of positive and negative outcomes which bias their processing of affective events. Normalising affective biases using computationally inspired interventions may represent a novel treatment approach for depression.

## Introduction

When learning about and interacting with the world, individuals vary in the extent to which their beliefs and behaviours are influenced by the events they experience. Often this variation displays an affective gradient with some individuals being more influenced by positive and others by negative events. For example, many people display an optimism bias, updating their beliefs to a greater extent following positive than negative outcomes (Sharot & Garrett, 2016). The opposite effect, a tendency to be more influenced by negative events, has been argued to cause illnesses such as depression (Mathews & MacLeod, 2005). Negative affective biases in depression have been reported using measures of attention (Gotlib, Krasnoperova, Yue, & Joormann, 2004), memory (Bradley, Mogg, & Williams, 1995; Nelson & Craighead, 1977) and learning (Eshel & Roiser, 2010). Consistent with their causal role in depression, interventions designed to target and reduce negative biases, such as cognitive behavioural therapy or more specific bias modification procedures can lead to improvement in symptoms (Browning, Holmes, Charles, Cowen, & Harmer, 2012; NICE, 2009). However, relatively little work has explored why individuals might develop negative biases in the first place. This question is of particular importance as understanding the mechanisms which lead to the development of affective bias is an essential first step in the development of novel treatments designed to alter this process and thus reduce symptoms of depression. One way of answering *why* individuals develop negative bias is to consider *when* negative biases might be the appropriate way to think about the world. In this study we draw on recent advances from the computational neuroscience of learning to investigate whether affective biases may be understood in terms of how informative an individual judges an event to be. Below we describe the conceptual framework of this proposal and then link this to the causal cognitive processes which underlie depression.

Recent computational work has demonstrated that individuals’ expectations are influenced more by those events which carry more information; that is, those events which improve predictions of future outcomes to a greater degree (T. E. J. Behrens, Woolrich, Walton, & Rushworth, 2007; Browning, Behrens, Jocham, O’Reilly, & Bishop, 2015; MacKay, 2003; Nassar et al., 2012). One factor which influences how informative an event is the changeability, or volatility, of the underlying association which is being learned. For example, imagine trying to learn what your colleagues think about your performance at work, based solely on their day-to-day feedback. One colleague seems to have a stable positive view of you, complimenting you on your work on 80% of the occasions you meet and never increasing or decreasing this frequency. In this case, each particular event (being complimented or not) provides little new information about what your colleague thinks about you, as you will always have an 80% chance of being complimented the next time you meet. In contrast, a second colleague’s appraisal of you seems to be more changeable, with periods when they think highly of you and compliment you regularly and others when they rarely compliment you at all. In this case each event provides more information; if you have recently been complimented by this colleague it is more likely that their opinion of you is currently high and they will compliment you the next time you meet (Figure 1B). When learning what your colleagues currently think about you, you should be more influenced by whether the second, more volatile, colleague compliments you or not, because this provides more useful information than the behaviour of the stable colleague.

Within a reinforcement learning framework, the influence of events on one’s belief is captured by the learning rate parameter, with a higher learning rate reflecting a greater influence of more recently experienced events (Sutton & Barto, 1998). Humans adjust their learning rate precisely as described above, using a higher learning rate for events, such as those occurring in a volatile context, which they estimate to be more informative (T. E. J. Behrens et al., 2007; Browning et al., 2015; Nassar et al., 2012). The neural mechanism by which this modification of learning rate is achieved is thought to depend on activity of the central norepinepheric system (Yu & Dayan, 2005), with increased phasic activity of the system, which may be estimated using pupilometry (Joshi, Li, Kalwani, & Gold, 2016), reporting the occurrence of more informative events (Browning et al., 2015; Nassar et al., 2012) and acting to enhance the processing of these events (Aston-Jones & Cohen, 2005).

This computational framework provides an overarching logic for when an individual might develop the negative affective biases which underlie depression; individuals should bias their processing towards negative events if they estimate that they are more informative than positive events. As well as providing a novel reformulation of why affective biases may develop, this framework also suggests a potential novel treatment target; that is, if a higher estimate of the information content of negative relative to positive events leads to negative affective bias and thus to symptoms of depression, interventions which redress these estimates this should reduce both negative biases and symptoms.

However, a number of critical questions concerning this account remain outstanding. Firstly, no previous study has demonstrated that humans maintain separate estimates of the information content of positive and negative events. We tested whether these estimates were maintained using a novel learning task (Figure 1) in which participant choice led to both positive and negative outcomes, with the volatility of the outcomes (and therefore their information content) being independently manipulated in separate task blocks. Secondly, for estimated information content to be a viable treatment target it must be malleable. We assessed this malleability by testing whether the volatility manipulation described above altered participants’ estimated information content, as reflected by the learning rates they used. Lastly, while activity of the central NE system has been argued to represent estimates of volatility, it is not clear whether or how this system might multiplex separate representations of the volatility of different classes of event, such as the positive and negative outcomes examined here. We investigate this using pupilometry as a measure of NE activity while participants completed the task. We hypothesised that humans maintain separable estimates of the information content of positive and negative outcomes, that we could measure and manipulate these estimates using our task and that phasic NE activity yoked to a specific type of outcome would track the volatility of that outcome.

## Methods and Materials

### Participants

30 English-speaking, individuals aged between 18 and 65 were recruited from the local community via advertisements. The number of participants recruited for the current cohort was selected to provide >95% power of detecting a similar effect size as that reported in a previous study in which a volatility manipulation was used to influence learning rate (Browning et al., 2015). Potential participants who were currently on a psychotropic medication or who had a history of neurological disorders were excluded from the study.

### General procedure

The study involved a single experimental session during which participants completed a novel learning task (described below) as well as standard questionnaire measures of depression (Quick Inventory of Depressive Symptoms, QIDS (Rush et al., 2003)) and anxiety (Spielberger State-Trait Anxiety Inventory, trait subscale, STAI (Spielberger, Gorsuch, & Lushene, 1983)) symptoms. The study was approved by the University of Oxford Central Research Ethics Committee. Written informed consent was obtained from all participants, in accordance with the Declaration of Helsinki.

### The Information Bias Learning Task (IBLT)

The information bias learning task (Figure 1) was adapted from a structurally similar learning task previously reported in the literature (T. E. J. Behrens et al., 2007; Browning et al., 2015). On each trial of the task participants were presented with two abstract shapes (letters selected from the Agathodaimon font) and chose the shape which they believed would result in the best outcome. On each trial one of the shapes, if chosen, would result in a win of 15p and one would result in a loss of 15p. These two outcomes were independent of each other so that a particular shape could be associated with one, both or neither of the win and loss outcomes (Figure 1C). As the two outcomes were independent participants had to separately learn the likely location of the win and the loss in the current trial. This learning was driven by the outcomes of previous trials and was used by participants to determine the most advantageous shape to choose on the current trial. Throughout the task the number and type of stimuli displayed during each phase of the trials was kept constant (Figure 1a) in order to minimise variations in luminance between trials.

**Figure 1.**
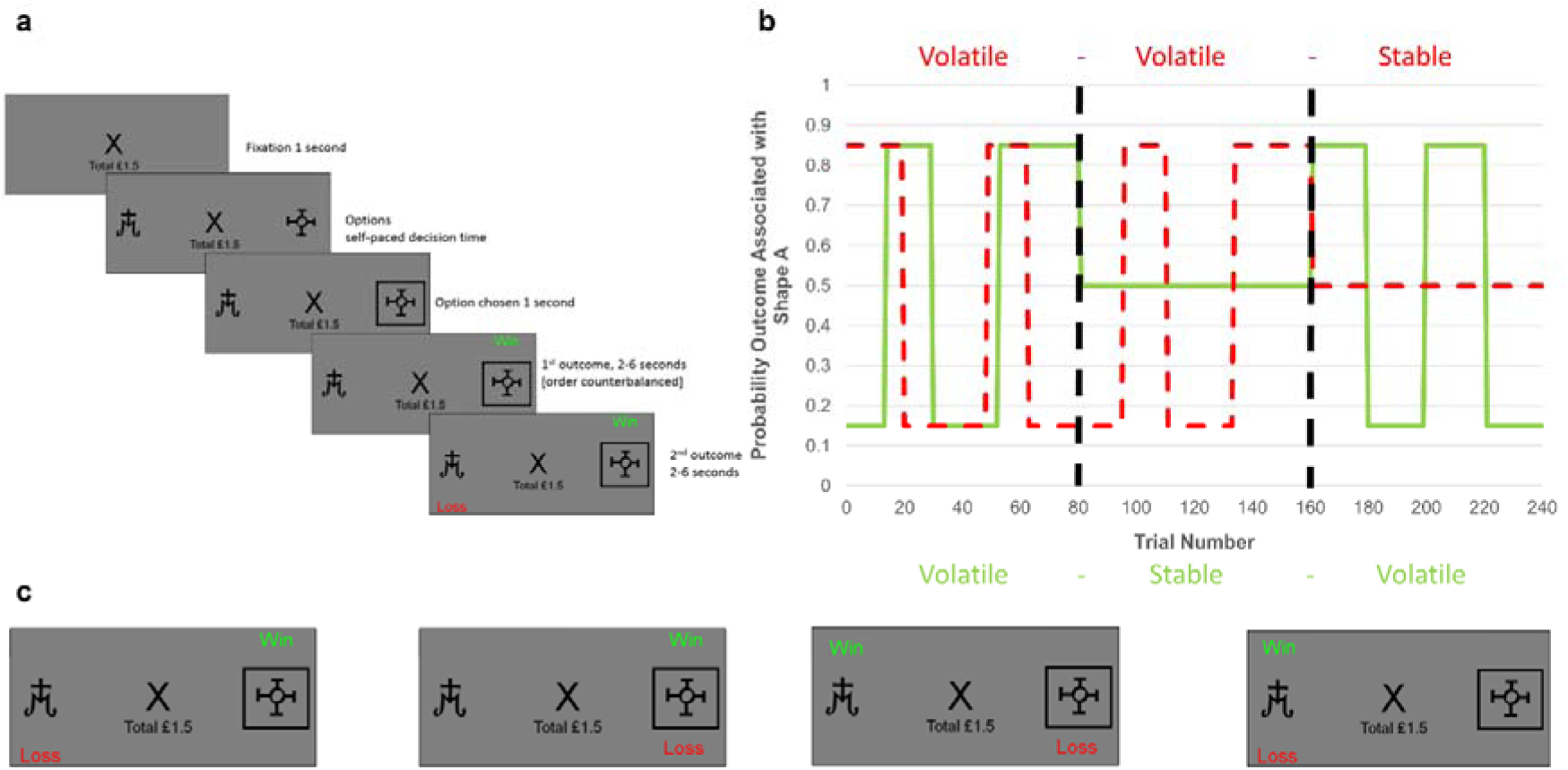
Task structure (A) Timeline of one trial from the learning task used in this study. Participants are presented with two shapes (referred to as shape “A” and “B”) and have to choose one. On each trial, one of the two shapes will be associated with a “win” outcome (resulting in a win of 15p) and one with a “loss” outcome (resulting in a loss of 15p). Using trial and error participants learn where the win and loss are likely to be found and use this information to guide their choice. (B) Overall task structure. The task consisted of 3 blocks of 80 trials each (i.e. vertical, dashed, dark lines separate the blocks). The y-axis represents the probability, *p*, that an outcome (win in solid green or loss in dashed red) will be found under shape “A” (the probability that it is under shape “B” is 1-*p*). The blocks differ in how volatile (changeable) the outcome probabilities are. Within the first block both win and loss outcomes were volatile, in the second two blocks one outcome was volatile and the other stable (here wins are stable in the second block and losses stable in the third block). The volatility of the outcome influences how informative that outcome is. Consider the second block in which the losses are volatile and the wins stable. Here, regardless of whether the win is found under shape “A” or shape “B” on a trial, it will have the same chance of being under each shape in the following trials, so the position of a win in this block provides little information about the outcome of future trials. In contrast, if a loss is found under shape “A”, it is more likely to occur under this shape in future trials than if it is found under shape “B”. Thus, for the second block losses provide more information than wins and participants are expected to learn more from them. (C) The four potential outcomes from a trial. Win and loss outcomes were independent, that is knowledge of the location of the win provided no information about the location of the loss. Because of this participants had to separately estimate where the win and where the loss would be on each trial in order to complete the task. This manipulation made it possible to independently manipulate the volatility of the two outcomes.

In total, the participants completed three blocks of 80 trials each, with a rest session between blocks. The same two shapes were used for all trials within a block, with different shapes being used between blocks. The outcome schedules were determined such that the probability that wins and losses were associated with shape A within a block always averaged 50%. In the volatile blocks the association between shape A and the outcome changed from 15 to 85% and back again in runs ranging from 14 to 30 trials. As described in the introduction, outcomes in the volatile blocks were more useful when predicting future outcomes, making them “informative”, whereas in the stable blocks outcome probabilities were fixed at 50%, making the outcomes “uninformative” in terms of predicting future trials (Figure 1B). In the first block of the task, both outcomes were volatile (informative), whereas in blocks 2 and 3 only one of the outcomes was volatile (informative) with the other being stable (uninformative). See supplementary materials for results from a control task in which volatility was kept constant, while the strength of the association between stimuli and outcomes (i.e. noise) was varied. The order in which blocks 2 and 3 were completed was counterbalanced across participants. Participants were paid all the money they had collected in the task, in addition to a £10 baseline payment. Choice data from the task was analysed by fitting a behavioural model which is described below. Alternative models are described and assessed in detail in the supplementary methods.

### Behavioural Model Used in Analysis of the IBLT

The primary measure of interest in the IBLT is the learning rate for wins and for losses in each of the three blocks. A simple behavioural model, based on that employed in related tasks (T. E. J. Behrens et al., 2007; Browning et al., 2015) was used to estimate learning rate. This model first estimated the separate probabilities that the win and loss would be associated with shape “A” using a Rescorla-Wagner learning rule (Rescorla & Wagner, 1972):

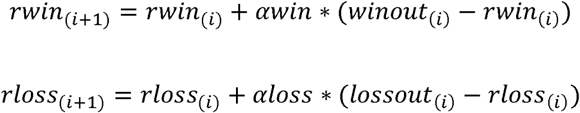
In these equations *rwin*_(_*_i_*_)_, which was initialised at 0.5, is the estimated probability that the win will be associated with shape “A” on trial i (NB the probability that the win is associated with shape “B” is 1*-rwin*_(_*_i_*_)_), *winout*_(_*_i_*_)_ is a variable coding for whether the win was associated with shape “A” (in which case the variable has a value of 1) or shape “B” (giving a value of 0) and *awin* is a free parameter, the learning rate for the wins, *rloss*_(_*_i_*_)_, *lossout*_(_*_i_*_)_ and *αloss* are the same variables for the loss outcome. These estimated outcome probabilities were then transformed into a single choice probability using a soft max function:

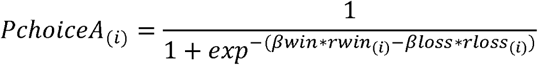
Where *PchoiceA*_(_*_i_*_)_ is the probability of choosing shape “A” on trial i, and *βwin* and *βloss* are inverse decision temperatures for wins and losses, respectively. The four free-parameters of this model (learning rates and inverse temperatures for wins and losses) were estimated separately for each task block and each participant by calculating the full joint posterior probability of the parameters, given participants’ choices, and then deriving the expected value of each parameter from their marginalised probability distributions (T. E. J. Behrens et al., 2007; Browning et al., 2015). Choice data from the first 10 trials of each block was not used when estimating the parameters as these trials were excluded from the pupil analysis (due to initial pupil adaption) (Browning et al., 2015; Nassar et al., 2012).

### Pupilometry Data

Full details of the preprocessing of the pupilometry data is provided in the supplementary methods. Preprocessing resulted in difference timeseries of pupil dilation data which represented the differential pupil dilation occurring during trials when the outcome (win or loss) was received relative to when it was not received over the six seconds after presentation of the outcomes. These timeseries were binned into 1 second bins to facilitate analysis.

### Data Analysis

Parameters derived from the computational models were transformed before analysis so that they were on the infinite real line (an inverse logit transform was used for learning rates and a log transform for inverse temperatures). Figures illustrate non-transformed parameters for ease of interpretation. The effect of the volatility manipulation on these transformed parameters was tested using a repeated measures ANOVA of data derived from the last two task blocks (i.e. when volatility was manipulated). In this ANOVA block volatility (win volatile block, loss volatile block) and parameter valence (wins, losses) were within subject factors and block order (win volatile first, loss volatile first) was a between subject factor. The critical term of this analysis is the block volatility x parameter valence interaction which tests for a differential effect of the volatility manipulation on the win and loss parameters.

The binned pupil timeseries data was analysed using a repeated measures ANOVA with time bin (1-6 seconds), block volatility (win volatile, loss volatile) and valence (wins, losses) as within subject factors and block order as a between subject factor. Again a block volatility x valence interaction tests for a differential effect of the volatility manipulation on the pupil dilation in response to wins vs. losses. In order to perform between subject correlations of the pupilometry data the mean relative dilation across the entire six second outcome period was also calculated for each participant and each block. In all analyses significant interactions were followed up by standard post-hoc tests.

## Results

Demographic details of the 30 participants are reported in Table 1.

**Table 1.**
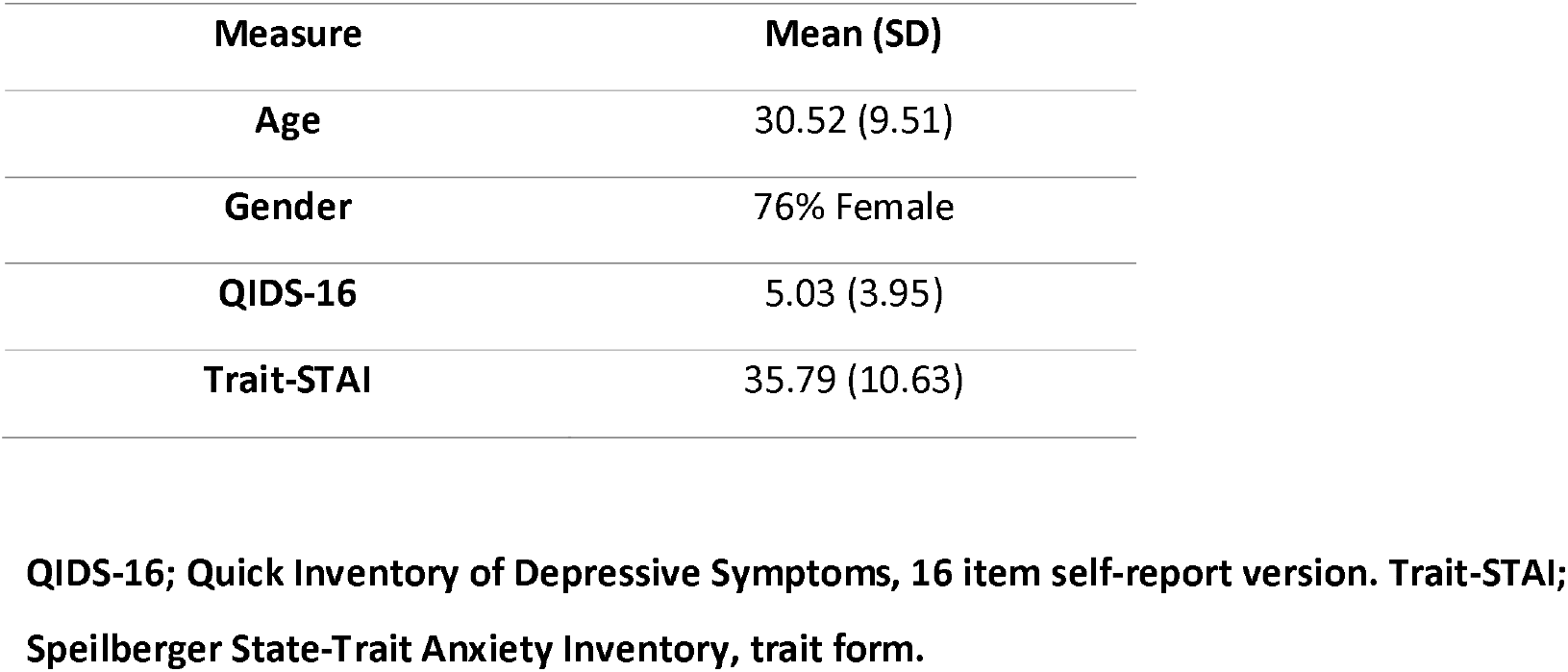
Demographic details of participants

| Measure | Mean (SD) |
| --- | --- |
| Age | 30.52 (9.51) |
| Gender | 76% Female |
| QIDS-16 | 5.03 (3.95) |
| Trait-STAI | 35.79 (10.63) |

**QIDS-16; Quick Inventory of Depressive Symptoms, 16 item self-report version. Trait-STAI; Speilberger State-Trait Anxiety Inventory, trait form.**

### Effect of Volatility Manipulation on Learning Parameters

As predicted, participants’ learning rates for positive and negative outcomes reflected the information content of the outcomes in the IBLT (block volatility x parameter valence; F(2,27) =26.488, *p* <0.001; Figure 2). Specifically, learning rates were higher for win (F(1,27) =16.59, *p* <0.001) and loss (F(1,27) =16.02, *p* <0.001) outcomes when they were volatile (informative) than when they were stable (not informative). Similarly the learning rate for wins was higher than that for losses when wins were more volatile than losses (F(1,27) =23.958, *p* <0.001) and the learning rate for losses was higher than for wins when losses were more volatile (F(1,27 =6.793, *p* <0.015). These results demonstrate that participants maintain independent estimates of the information content of positive and negative outcomes and that it is possible to alter these estimates using a simple volatility manipulation. In contrast to the effects on learning rate there were no significant effects of the task on the inverse temperature parameter of the learning model (F(1,27) =0.038, *p*=0.846) indicating that, as intended, the volatility manipulation specifically altered learning rate rather than the relative weights placed on positive and negative outcomes (Huys, Pizzagalli, Bogdan, & Dayan, 2013). See the Supplementary Materials for additional analysis of the behavioural results as well as an additional control experiment.

**Figure 2.**
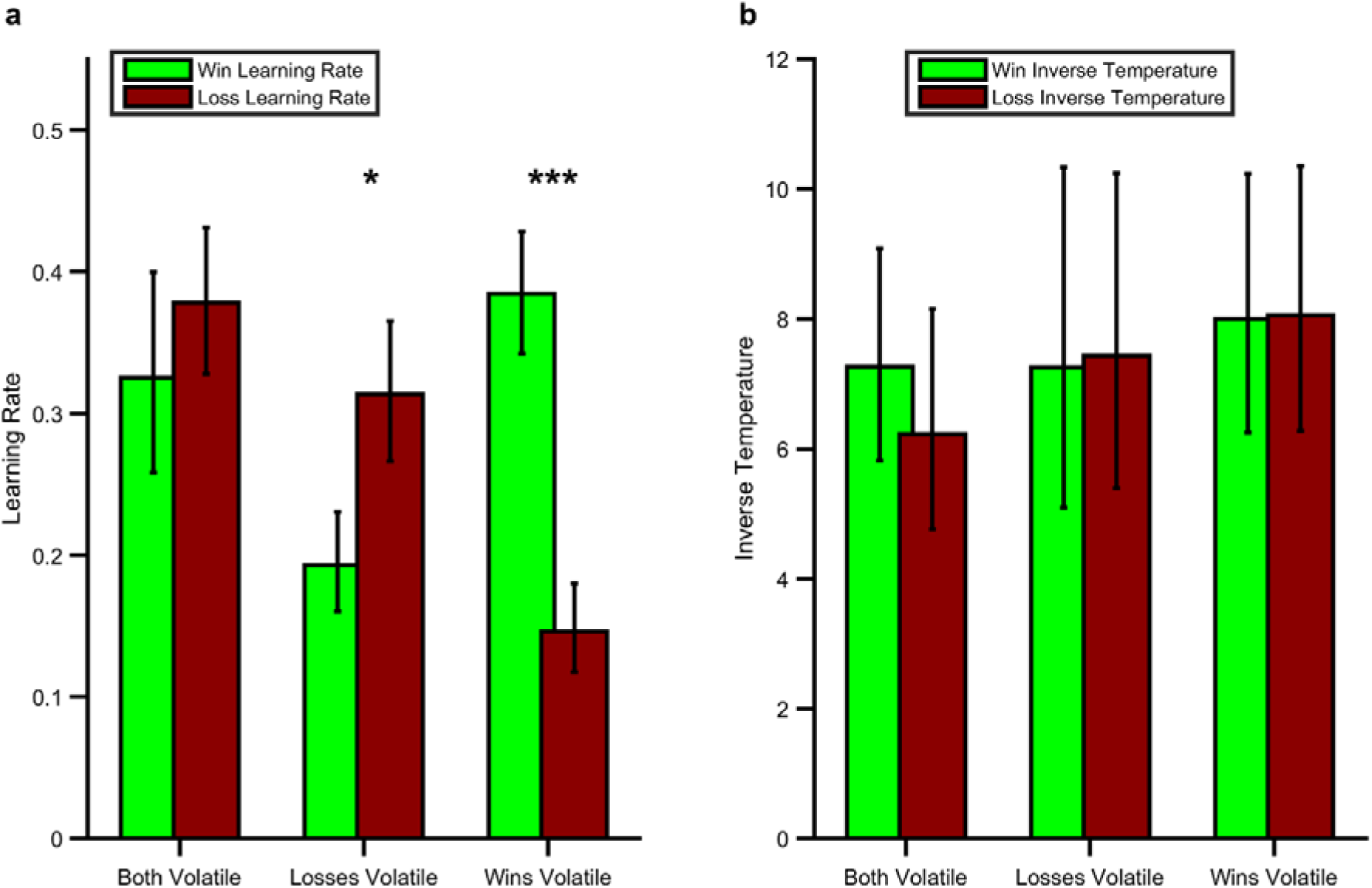
Effect of Volatility Manipulation on Participant Behaviour. (A) Mean (SEM) learning rates for each block of the IBLT. As can be seen the win learning rates (light green bars) and loss learning rate (dark red bars) varied independently as a function of the volatility of the relevant outcome (F(1,27)=26.488, p<0.001), with a higher learning rate being used when the outcome was volatile than stable (*p<0.05, *** p<0.001 for pairwise comparisons). (B) No effect of volatility was observed for the inverse temperature parameters (F(1,27)=0.038, p=0.846).

### Effect of Volatility Manipulation on Pupil Dilation

Next, we investigated the extent to which central NE activity, as estimated using pupilometry, was related to the information content of positive and negative outcomes in the IBLT. Consistent with the behavioural findings a significant interaction between block volatility and outcome valence was found for the degree to which participants’ pupils dilated in response to outcome receipt (Figure 3; F(1,27)=4.9; p=0.04). In other words, participants’ pupils dilated more on receipt of an outcome when that outcome was volatile (informative) than when it was stable (not informative). This effect was not further modified by the time bin following outcome (block volatility x outcome valence x time; F(5,135)=0.340, p=0.565). Analysing the positive and negative outcomes separately indicated that the effect of block volatility was significant for the loss outcomes (F(1,27)=7.597, p = 0.01), but not for the win outcomes (F(1,27)=0.157, p = 0.695).

**Figure 3.**
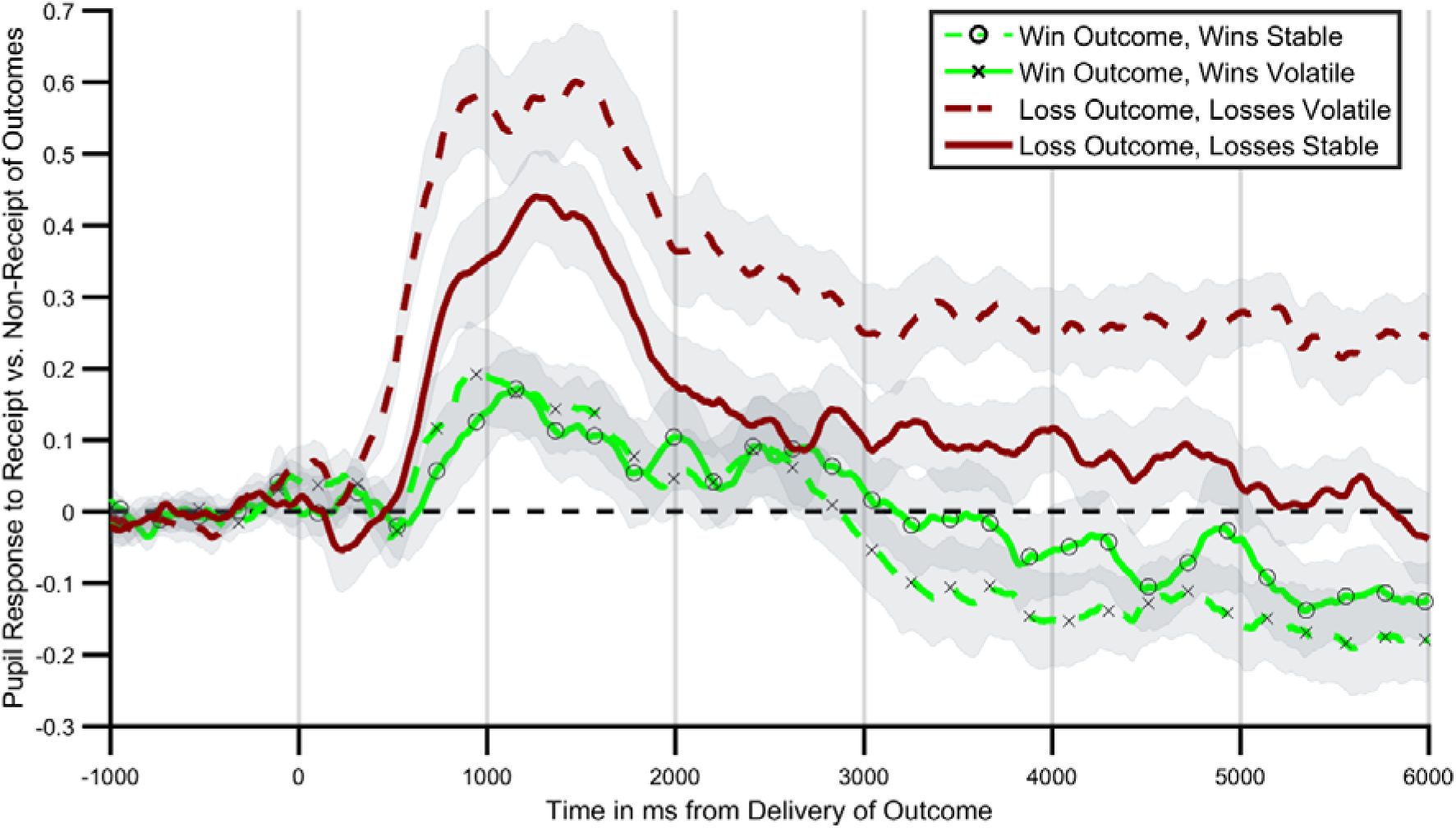
Pupil response to outcome delivery during the IBLT. Lines illustrate the mean pupil dilation to the receipt relative to non-receipt of an outcome across the 6 seconds after outcomes are presented. Light green lines (with crosses and circles) report response to win outcomes, dark red lines report response to loss outcomes. Solid lines report blocks in which the wins were more informative (volatile), dashed lines blocks in which losses were more informative. As can be seen pupils dilated more when the relevant outcome was more informative, with this effect being particularly marked for loss outcomes. Shaded regions represent the SEM.

### Relationship Between Choice Behaviour and Pupil Dilation

As central NE activity is thought to mediate the effect of outcome information content on participant choice (Yu & Dayan, 2005), there should be a relationship between how much a participant’s pupils differentially dilate in response to an outcome during the informative and non-informative blocks and the degree to which that participant adjusts their learning rate between blocks for the same outcome. We tested this by assessing the correlation between the change in mean pupil response between blocks and the change in behaviourally estimated learning rates, separately for wins and losses. As can be seen (Figure 4) the change in pupil response to loss outcomes between blocks was significantly correlated with the change in loss learning rate (r(28)=0.5, p=0.009) but pupil response to win outcomes was not correlated with change in win learning rate (r(28)=−0.08, p=0.7).

**Figure 4.**
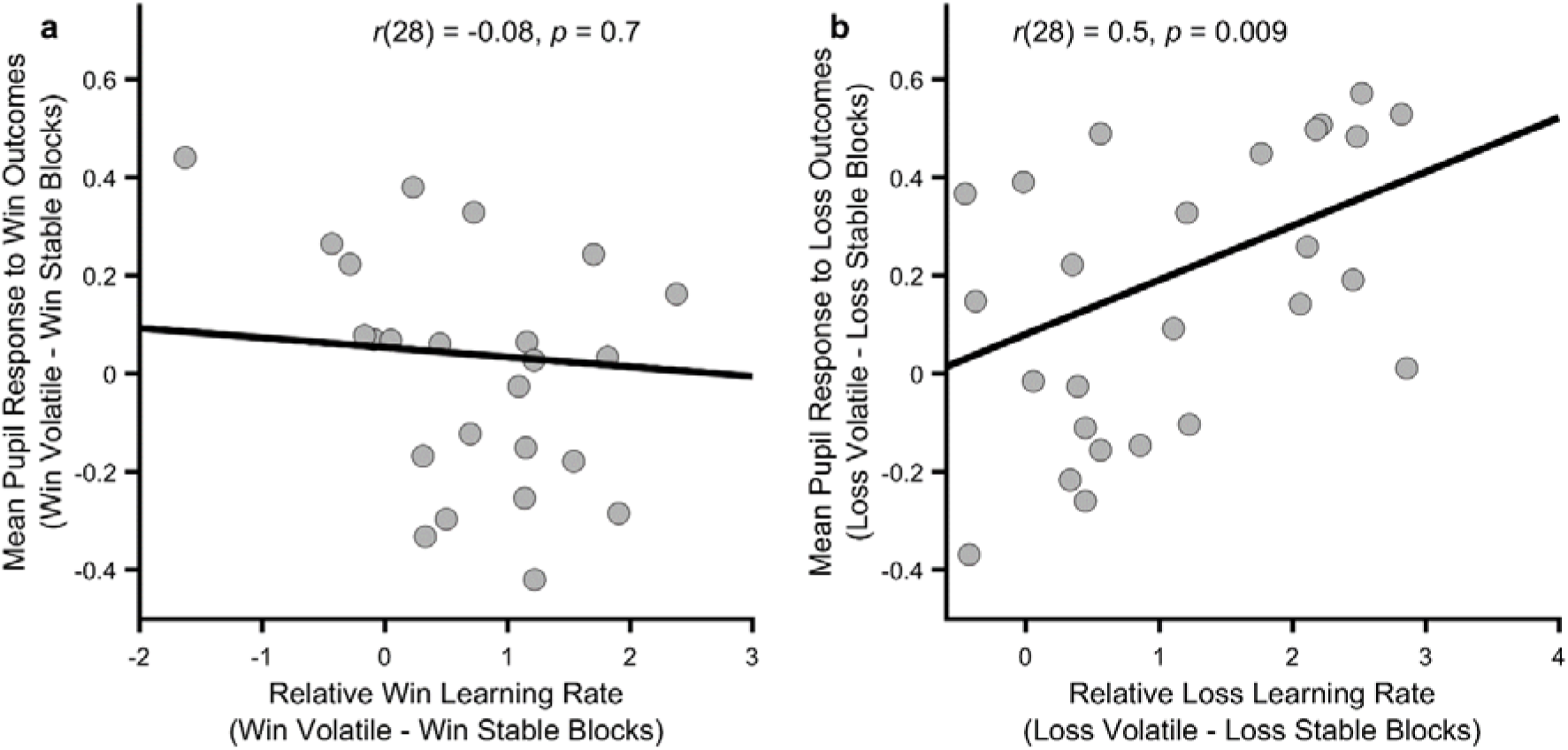
Relationship between behavioural and physiological measures. The more an individual altered their loss learning rate between blocks, the more that individual’s pupil dilation in response to loss outcomes differed between the blocks (panel b; p=0.009), however no such relationship was observed for the win outcomes (panel a; p=0.7). Note that learning rates are transformed onto the real line using an inverse logit transform before their difference is calculated and thus the difference score may be greater than ±1.

## Discussion

Humans adapt the degree to which they are influenced by positive and negative outcomes in response to how informative they estimate those outcomes to be. These estimates produce an affective bias in learning, with a higher learning rate being used for the class of outcome which is most informative, and are malleable. They may thus represent a novel, computationally defined cognitive treatment target for depression. A physiological measure of central NE activity was associated with this process, although this was only seen convincingly for loss outcomes.

Previous work has demonstrated that humans adapt their learning in response to subtle statistical aspects of the environment, such as employing an increased learning rate in volatile, or changeable, contexts (T. E. J. Behrens et al., 2007; Browning et al., 2015; Nassar et al., 2012). This suggests that learners maintain an estimate of how useful, or informative, an event is and learn more from events they estimate to be more informative. The current study extends this work by providing evidence that humans are able to maintain independent estimates of the information content of different classes of event, in this case positive and negative outcomes (winning vs. losing money). The parallel representation of estimated information content of wins and losses provides a mechanism by which individuals may come to be generally more influenced by events of one class than another. In the case of depression, patients have been shown to be more influenced by negative events, for example tending to remember more negative than positive events (Bradley et al., 1995), attend to negative more than positive events (Gotlib et al., 2004) and learn more from negative and less from positive outcomes (Eshel & Roiser, 2010). The results of the current study suggest that these observed negative biases may all be understood as a consequence of patients estimating that the information content of negative relative to positive events was higher than non-patients. As the negative biases described above are believed to be causally related to symptoms of depression (Mathews & MacLeod, 2005), and interventions designed to alter negative biases can reduce symptoms (Browning et al., 2012; NICE, 2009), these results raise the possibility that novel interventions which target expected information content may act to reduce symptoms of the illness. Of course, identifying potential targets for treatment and showing that they may be altered experimentally as done in this paper is only the first step in the development of new treatments. The next step, analogous to a phase 2a study in drug development (Ciociola et al., 2014), is to assess the initial efficacy of a potential intervention which engages the target in a clinical population. A study designed to do this is currently underway using the volatility manipulation described in this paper (study identifier NCT02913898).

In the current study we investigated the link between the learning rate used by participants, which provides a behavioural index of how informative they estimate an outcome to be, and pupil dilation which has been shown to correlate with central norepinepheric activity (Joshi et al., 2016). Pupil dilation in response to outcome receipt differed as a function of the information content of the outcome, although this was only significant for losses. Specifically, when losses were informative, the difference in pupil dilation between trials in which a loss was received and when it was not received was greater than when the losses were not informative. This result is similar to previously reported findings of an increased pupil response to outcomes in a volatile context (Browning et al., 2015; Nassar et al., 2012), although these earlier studies reported a general increase in pupil dilation rather than a dilation conditioned on receipt of the outcome. A possible explanation of this difference is that, in the current study, one of the outcomes (win or loss) was always volatile and presentation order of the outcomes was randomised. Therefore, in contrast to the previous studies in which only one class of outcome was used, the relevant volatility signal required to perform the task in the current study was dependent on the outcome presented. In other words, the volatility signal found in the pupil data from the current study is of the form required for participants to accurately perform the task. This suggests a degree of flexibility of the pupillary volatility signal, in that it may reflect the general volatility of a learned association or the volatility of specific dimensions of more complex associations depending on task demands. It is not clear whether these general and specific volatility signals are produced by a single or separate neural systems, although it may be possible to address this question using a task in which the total volatility of all task outcomes is manipulated independently of the volatility of the individual outcomes. The finding that the volatility signal in the current task modifies pupillary response to outcome receipt may also explain why the pupilometry measure was sensitive only to loss and not win outcomes; receipt of a loss lead to a greater pupil dilation overall than a win (see Sup Figure 6) and thus the effect of estimated outcome information, which modifies the relative dilation observed when an outcome is received, may be less apparent for wins.

The pupilometry measure included in the current study raises the possibility that estimated information content may be influenced by pharmacological as well as cognitive interventions. Pupil size is influenced by the activity of a number of central neurotransmitters including norepinephrine (Joshi et al., 2016) and previous work exploring the neural systems which control response to volatility have predicted a key role for NE (Yu & Dayan, 2005) suggesting it as an obvious pharmacological target. A single study has reported an effect of atomoxetine, a norepinephrine reuptake inhibitor, on learning in a volatile environment (Jepma et al., 2016) although no previous work has examined the effect of a pharmacological intervention on learning to positive vs. negative outcomes. It would be interesting to test whether a pharmacological manipulation of norepinepheric function was able to modify the outcome specific volatility effect demonstrated in this paper as such an effect may indicate a clinically useful interaction between pharmacological and cognitive interventions.

The information content of an outcome is not solely a function of the volatility of its occurrence. Other factors, such as the strength of the association between a stimulus, or action, and the subsequent outcome, sometimes called the “expected uncertainty” (Yu & Dayan, 2005) of the association, will also influence how informative the outcome is. Outcomes in the IBLT task reported in this paper vary in terms of both volatility and expected uncertainty, with both of these factors predicted to influence learning rate in the same direction (i.e. both factors should increase learning rate in the volatile blocks). A control experiment (see supplementary materials) in which volatility was kept constant but expected uncertainty varied found no effect on learning rate suggesting that the current findings were likely to be due to the effects of volatility rather than expected uncertainty. However, it would be interesting in future studies to explore whether it was possible to use manipulations of expected uncertainty, in the same way that volatility is used in this study, to induce a preference for positive over negative events. This may provide an alternative approach to engaging and altering expected information content than the volatility based effect reported here.

The current study demonstrates that human learners maintain separable estimates of the information content of positive and negative outcomes and provides an initial proof of principle as to how these estimates may be modified. The study illustrates a little explored application of computational techniques in cognitive neuroscience; they may be used to identify novel treatment targets and by so doing spur the development of new and more effective treatments.

## Supplementary Methods

### Further Details of the IBLT

The task was presented on a VGA monitor connected to a laptop computer running Presentation software version 18.3 (Neurobehavioural Systems). Participants’ heads were stabilised using a head-and-chin rest placed 70 cm from the screen on which an eye tracking system was mounted (Eyelink 1000 Plus; SR Research). The eye tracking device was configured to record the coordinates of both of the eyes and pupil area at a rate of 500 Hz. The abstract shapes of the learning task were drawn on either side of a fixation cross which marked the middle of the screen and were offset by around 7° visual angle. The two outcomes (win and loss) were displayed on the screen in randomised order for a jittered interval of 2-6 (mean 4) seconds. Auditory stimuli lasting 0.7 seconds were played when participants received a win (“chi-ching” sound) or loss (error buzz). Participants’ accumulated total winnings was displayed under the fixation cross and was updated based at the beginning of the subsequent trial.

### Preprocessing of Pupil Data

Blinks were identified using the Eyelink system’s built in filter and were then removed from the data. Missing data points (including blinks) were linearly interpolated. The resulting trace was subjected to a low pass Butterworth filter with a cut-off of 3.75 Hz and then z transformed across the session (Browning et al., 2015; Nassar et al., 2012). The pupil response to the win and the loss outcomes were extracted separately from each trial, using a time window based on the presentation of the outcomes. This included a 1-s baseline period before the presentation of the outcome, and a 6-s period following outcome presentation. Baseline correction was performed by subtracting the mean pupil size during the 1 second baseline period prior to the presentation of each outcome, from each time point in the post outcome period. Individual trials were excluded from the pupilometry analysis if more than 50% of the data from the outcome period had been interpolated (mean =7% of trials) (Browning et al., 2015). One participant was excluded from the pupilometry analysis as more than 99% of their trials were excluded on this basis. The first 10 trials from each block were not used in the analysis as initial pupil adaption can occur in response to luminance changes in this period (Browning et al., 2015; Nassar et al., 2012). The preprocessing resulted in two sets of timeseries per participant, one set containing pupil dilation data for each included trial when the win outcomes were displayed and the other when the loss outcomes were displayed. A difference timeseries, calculated as the mean pupil response to the receipt vs. non-receipt of the outcome in each block was then calculated which allowed for assessment of how the volatility of a specific outcome influenced dilation in response to receiving vs. not receiving that outcome (See below for a complementary regression analysis of this data). In order to statistically compare these timeseries the mean of each 1 second time bin after outcome presentation was calculated.

### Alternative Behavioural Models and Model Selection

The behavioural model used in this study (Referred to as model 1 below) was developed based on the models used in previous studies in which volatility is manipulated (T. E. Behrens, Hunt, Woolrich, & Rushworth, 2008; T. E. J. Behrens et al., 2007; Browning et al., 2015) and to allow for the possibility that differential behaviour in response to win and loss outcomes may have arisen due to changes in learning rate (captured using separate win and loss learning rates) or outcome sensitivity (captured using separate inverse temperature parameters). However, it is possible that this model does not provide the best fit to participant choice data. In order to assess this possibility we compared the fit of this model against a range of comparator models using the Bayesian Information Criteria (BIC) metric, which includes a penalty term for model complexity.

Model 2: It is possible for participants to perform our task without learning the independent probability of the win and loss outcomes, but rather by taking a model-free (Daw, Gershman, Seymour, Dayan, & Dolan, 2011) approach in which the overall value of each shape was learned.

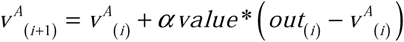
Here the value of shape A (*V^A^*) initiates at 0 on trial 1, and is updated on every trial based on the joint outcome (i.e. the win – loss for that shape) of the trial (*out*_(_*_i_*_)_), which can be −1, 0 or 1 with a single learning rate (*αvalue*). The estimated relative values of the 2 shapes were then transformed into a choice probability using a softmax function with a single inverse temperature parameter.

Model 3: An alternative approach, described by Behrens and colleagues (T. E. J. Behrens et al., 2007) estimates trialwise volatility within a fully Bayesian framework. For this model we used Behrens’ Bayesian learner to independently estimate the expected probabilities of the win and loss outcomes during the task (note that there are no free parameters for this learner). These estimates were then combined using the same selector model described in the main text with two inverse temperature parameters.

Model 4: This was a slightly simpler version of Model 1 in that it employed only a single inverse temperature parameter allowing assessment of the degree to which using 2 such parameters influenced model fit.

Model 5: Finally, we tested a slightly more complex version of Model 4 by including a risk parameter *γ* as used in previous studies, which modulates the estimated probabilities of wins and losses in a non-linear way. Risk parameters have been shown to account for non-normative aspects of human choice, particularly when outcome probabilities are particularly high or low:

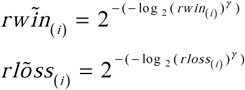
A summary of the five models can be found in Supplementary Table 1 below:

**Table S1:** **Description of Comparator Models**

| Model Name | Number of Learning Rate Parameters | Number of Inverse Temperature Parameters | Notes |
| --- | --- | --- | --- |
| 1. | 2 | 2 | Model used in paper |
| 2. | 1 | 1 | Model-free learner |
| 3. | 0 | 2 | Bayesian learner |
| 4. | 2 | 1 | Single inverse temperature model |
| 5. | 2 | 1 | Additional risk parameter |

All models were fitted to participant data using the same procedure described in the main paper. BIC scores for each model are illustrated in figure S1 below (note that lower scores indicate a better fit). As can be seen the model reported in the main paper (Model 1) fits the data best. The single inverse temperature model (Model 4) performs almost as well, with the other models performing less well.

**Supplementary Figure S1:**
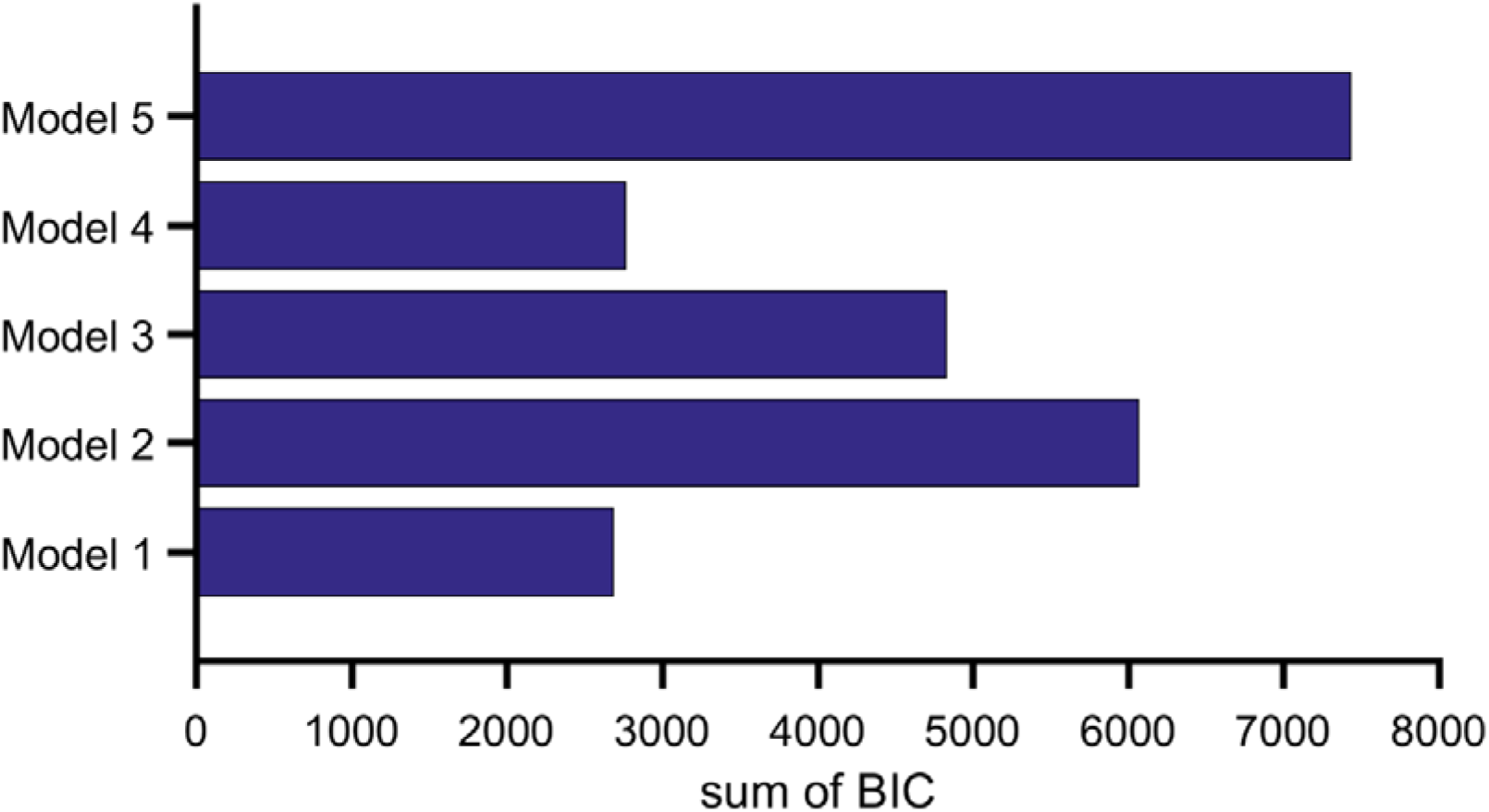
**BIC Scores for Comparator Models (see table S1 for model descriptions). Smaller BIC scores indicate a better model fit.**

## Supplementary Results

### Switch-Stay Analysis of Behaviour

The IBLT includes both positive and negative outcomes which are independent of each other. As a result the task contains trials in which both positive and negative outcome encourage the same behaviour in future trials (e.g. when the win is associated with shape A and the loss with shape B, both outcomes encourage selection of shape A in the following trial) as well as trials in which the positive and negative outcomes act in opposition (e.g. when both outcomes are associated with shape *A*, then the win outcome encourages selection of shape A in the next trial and the loss outcome encourages selection of shape B). This second type of trial provides a simple and sensitive means of assessing how the volatility manipulations alters the impact of win and loss outcomes on choice behaviour in the task blocks. Specifically an increased influence of win outcomes (e.g. when wins are volatile) should lead to:

a. A decreased tendency to change (shift) choice when both win and loss outcomes are associated with the chosen shape in the current trial
b. An increased tendency to change (shift) choice when both win and loss outcomes are associated with the unchosen shape in the current trial.

This analysis does not depend on any formal model and thus can be used to complement the model based analysis reported in the main paper. We calculated the proportion of shift trials separately for trials in which both outcomes were associated with the chosen or unchosen shape for each of the three blocks. Consistent with the model based analysis, participants switched significantly less frequently when both outcomes were associated with the chosen option in the win relative to loss informative blocks (Figure S2; F(1,27)=6.193, p=0.019) and switched significantly more frequently when both outcomes were associated with the unchosen option in the win relative to loss informative blocks (Figure S2; F(1,27)=4.353, p=0.047). This indicates that the results reported in the main paper are unlikely to be dependent on the exact form of the behavioural model used to derive the learning rate parameter.

**Supplementary Figure S2:**
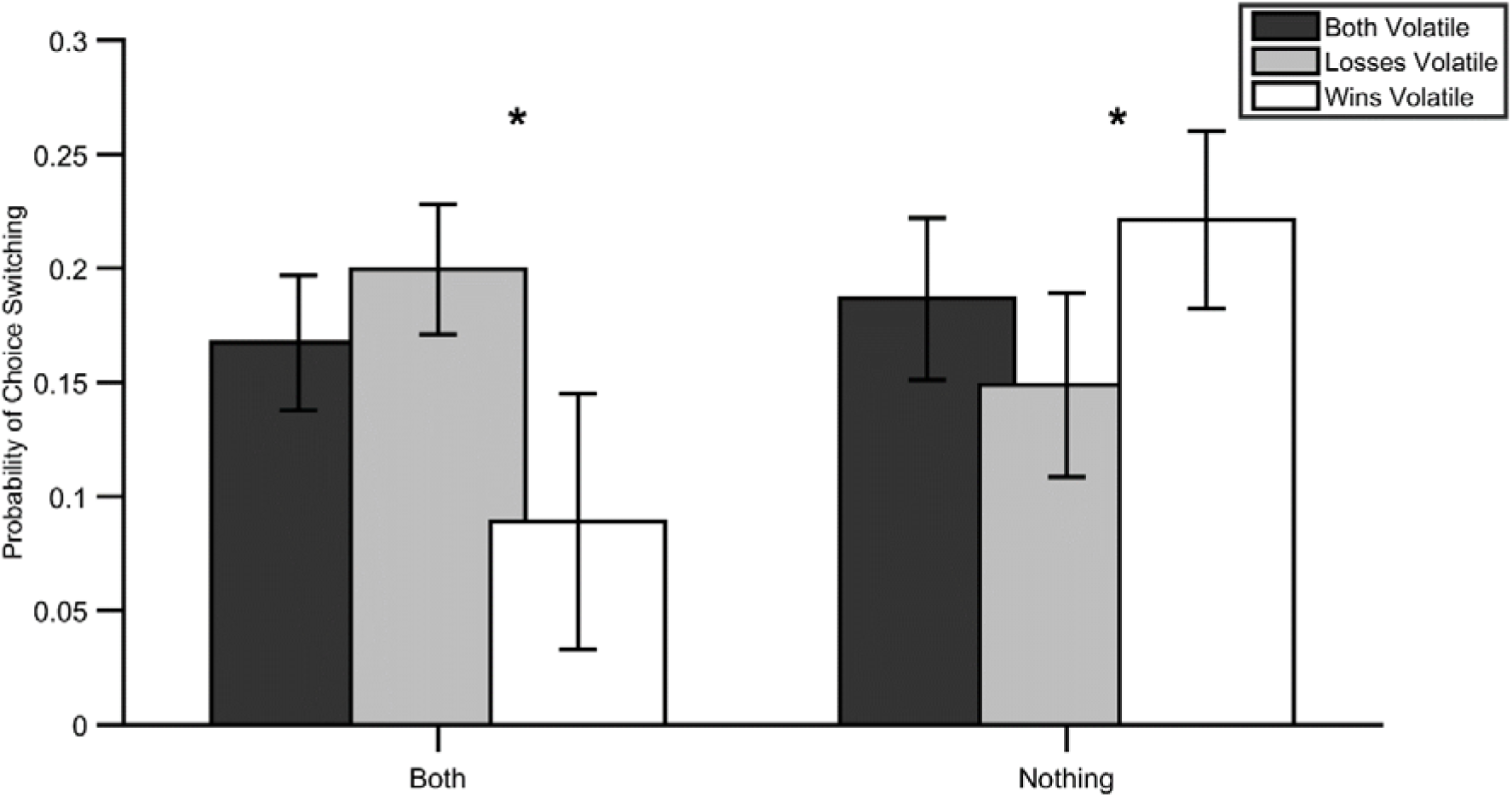
Analysis of switching behaviour in the IBLT task. The mean (SEM) probability of switching choice in the subsequent trial is plotted separately for trials in which both win and loss outcome are associated with the chosen option (“both”) and the non-chosen option (“nothing”). The columns represent the probability of switching in the first block of the task when both outcomes were informative/volatile (dark columns), in the block in which losses were more informative (grey columns) and the block in which wins were more informative (white column). As can be seen, when wins are more informative than losses (i.e. white bars), participant choice is more influenced by the win relative to loss outcomes than when losses are more informative (grey bars). Specifically, participants are more likely to stick with a choice which has just resulted in both a win and a loss and are more likely to switch to a choice if they didn’t choose it when the wins are informative. *=*p*<0.05 for comparison between win informative and loss informative blocks.

### Expected vs Unexpected Uncertainty

When learning, a number of different forms of uncertainty can influence behaviour. One form, which is sometimes called “unexpected uncertainty” (Yu & Dayan, 2005) is caused by changes in the associations being learned (i.e. volatility) and is the main focus of this paper (see main text for a description of how volatility influences learning). A second form of uncertainty, sometimes called “expected uncertainty”(Yu & Dayan, 2005) arises when an association between a stimulus or action and the subsequent outcome is more or less predictive. For example, this form of uncertainty is lower if an outcome occurs on 90% of the times an action is taken and higher if the outcome occurs on 50% of the time an action is taken. Normatively, expected uncertainty should influence learning rate—a less predictive association (i.e. higher expected uncertainty) leads to more random outcomes which tell us less about the underlying association we are trying to learn, so learners should employ a lower learning rate when expected uncertainty is higher. In the task described in this paper both the expected and unexpected uncertainty differ between blocks. Specifically, when an outcome is stable in the task it occurs on 50% of trials, whereas when it is volatile it varies between occurring on 85/15% of trials. Thus the stable outcome is, at any one time, also less predictable (i.e. noisier) than the volatile outcome. This task schedule was used as a probability of 50% for the stable outcome improves the ability of the task to accurately estimate learning rates (it allows more frequent switches in choice). Further both forms of uncertainty would be expected to reduce learning rate in the stable blocks and increase it in the volatile block of the task. However, this aspect of the task raises the possibility that the observed effects on behaviour described in the main paper may arise secondary to differences in expected uncertainty (noise) rather than the unexpected uncertainty (volatility) manipulation. In order to test this possibility we developed a similar learning task in which volatility was kept constant and expected uncertainty was varied (Figure S4). In this task, participants again had to choose between two shapes in order to win as much money as possible, however on each trial 100 “win points” and 100 “loss points” were divided between the two shapes and participants received money proportional to the number of win points – loss points of their chosen option. Thus, a win and loss outcome occurred on every trial of this task, but the magnitude of these outcomes varied. During the task, participants had to learn the expected magnitude of wins and losses for the shapes rather than the probability of their occurrence. This design (Figure S4a) allowed us present participants with schedules in which the volatility (i.e. unexpected uncertainty) of win and loss magnitudes was constant but the noise (expected uncertainty) varied (Figure S4b). Otherwise the task was structurally identical to the IBLT with 240 trials split into 3 blocks. We recruited a separate cohort of 30 healthy participants who completed this task and then estimated their learning rate using a model which was structurally identical (i.e. 2 learning rates and 2 inverse temperature parameters) to that used in the main paper (Model 1). As can be seen (Figure S4c), there was no effect of expected uncertainty on participant learning rate (block information x parameter valence; F(1,28)=1.97, p=0.17) during this task. This suggests that the learning rate effect seen in the IBLT cannot be accounted for by differences in expected uncertainty and therefore is likely to have arisen due to the unexpected uncertainty (volatility) manipulation. Inverse decision temperature did differ between block (F(1,28)=5.56, p=0.026). As can be seen in Figure S4d, there was a significantly higher win inverse temperature during the block in which the losses had lower noise (F(1,28)=9.26,p=0.005) and when compared to the win inverse temperature when wins had lower noise (F(1,28)=5.35,p=0.028), but no equivalent effect for loss inverse temperature. These results suggest that, if anything, participants were more influenced by noisy outcomes.

**Figure S4:**
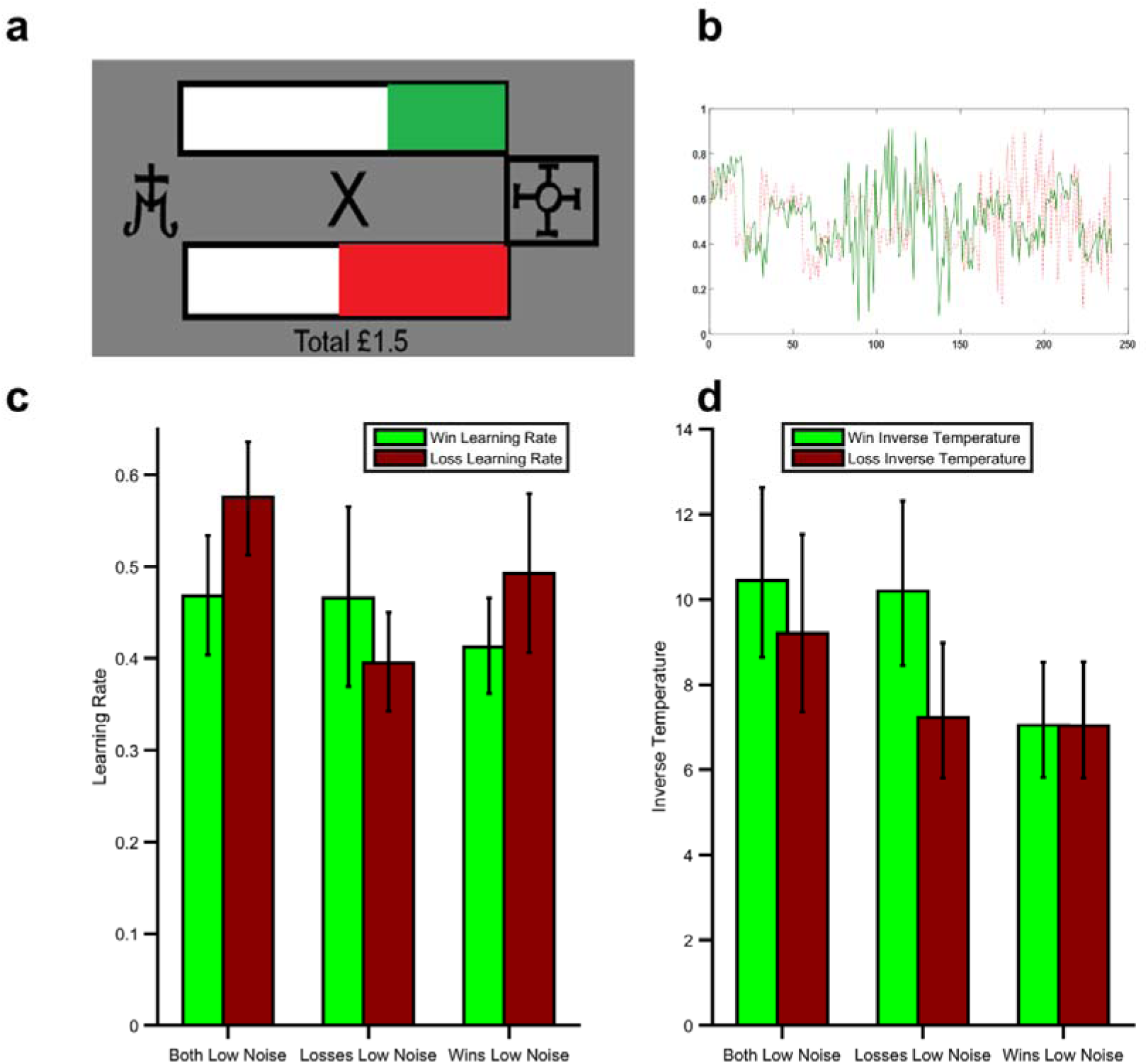
Magnitude Task. A) example outcome screen from the task. Participants chose between two shapes. Each shape, if chosen, resulted in winning a proportion of 100 win points (bar on top of fixation cross with green fill) and loosing a proportion of 100 loss points (bar under fixation cross with red fill), with participants receiving the differnce between the two. B) The task schedule for win (green) and loss (red) magnitdues included 3 blocks; in the first block both outcomes had low expected uncertainty (noise), in the last two blocks one outcome had high and the other low expected uncertianty. The volatility of the outcomes was constant across blocks. C) Participants did not significantly adjust their learning rates in response to expected uncertainty and D) inverse temperature for wins was increased during the block in which the losses had lower noise, with no effect on loss inverse temperature.

### Regression Analysis of Pupil Data

The analysis of pupil data reported in the main text examines the effect of block information content (i.e. win volatile vs. loss volatile) and outcome receipt on the pupil response to win and loss outcomes. However a number of other factors may also influence pupil dilation such as the order in which the outcomes were presented and the surprise associated with the outcome (Browning et al., 2015). In order to ensure that these additional factors could not account for our findings we ran a regression analysis of the pupil data from the IBLT task. In this analysis we derived, for each participant, trialwise estimates of the outcome volatility and outcome surprise of the chosen option using the Ideal Bayesian Observer reported by Behrens et al. (T. E. J. Behrens et al., 2007). These estimates were entered as explanatory variables alongside variables coding for outcome order (i.e. win displayed first or second), outcome of the trial (outcome received or not) and an additional term coding for the interaction between the outcome volatility and outcome of the trial (i.e. analogous to the pupil effect reported in Figure 3 of the main paper). Separate regression analyses were run for each 2ms timepoint across the outcome period, for win and loss outcomes and for each participant. This resulted in timeseries of beta weights representing the impact of each explanatory factor, for each participant and for win and loss outcomes. As can be seen in Figure S5 below, consistent with the results reported in the paper this analysis revealed a significant volatility x outcome interaction for loss outcomes (F(1,27)=6.249, p = 0.019), with no effect for wins (F(1,27)=0.215, p = 0.646). This result indicates that pupil effects reported in the main paper are not the result of outcome order or surprise effects on pupil dilation.

**Supplementary Figure S5.**
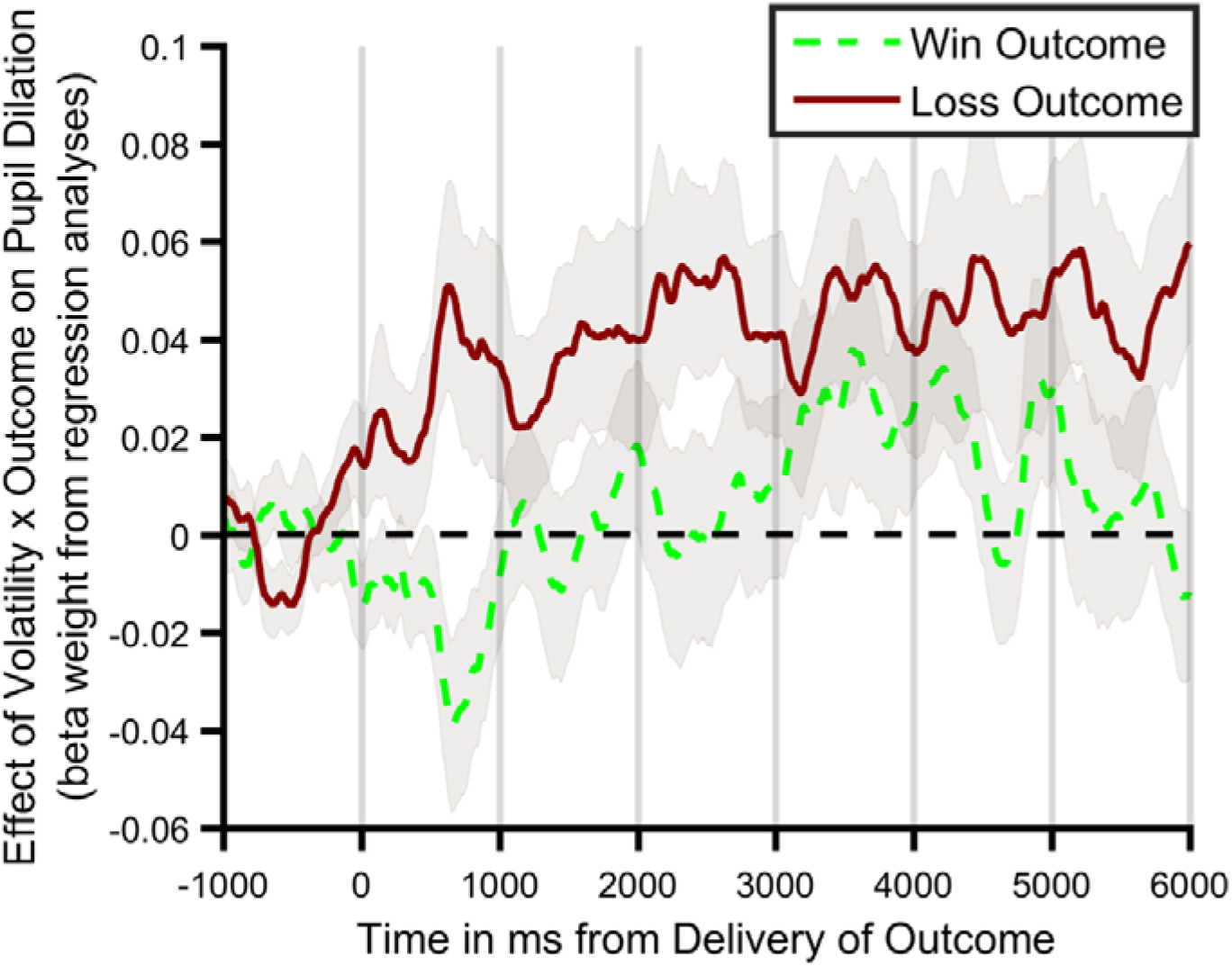
Regression analysis of pupil data. The mean (SEM) beta weight of the volatility x outcome regressor of the regression analysis of the pupil data is shown separately for win (green) and loss (red) outcomes. The loss regressor differs significantly from 0 for the loss outcomes indicating that, across participants, pupil dilation was greater in response to an outcome in the volatile than stable block for losses. No significant effect was observed for win outcomes.

### Post-Hoc Analysis of Pupil Data

Figure 3 from the paper illustrates the difference in pupil dilation between trials in which an outcome was received and those in which the outcome was not received. In order to further investigate this effect Figure S6 below separately plots the mean pupil response for trials in which the outcome was and was not received. As can be seen, whereas there is relatively little difference in pupil response during the win trials, there is a large difference in dilation between trials on which a loss is received and those in which no loss is received. Further, the effect of loss volatility is seen to both increase dilation on receipt of a loss and reduce dilation when no loss is received, suggesting that the effect of the volatility manipulation is to exaggerate the effect of the outcome.

**Supplementary Figure S6.**
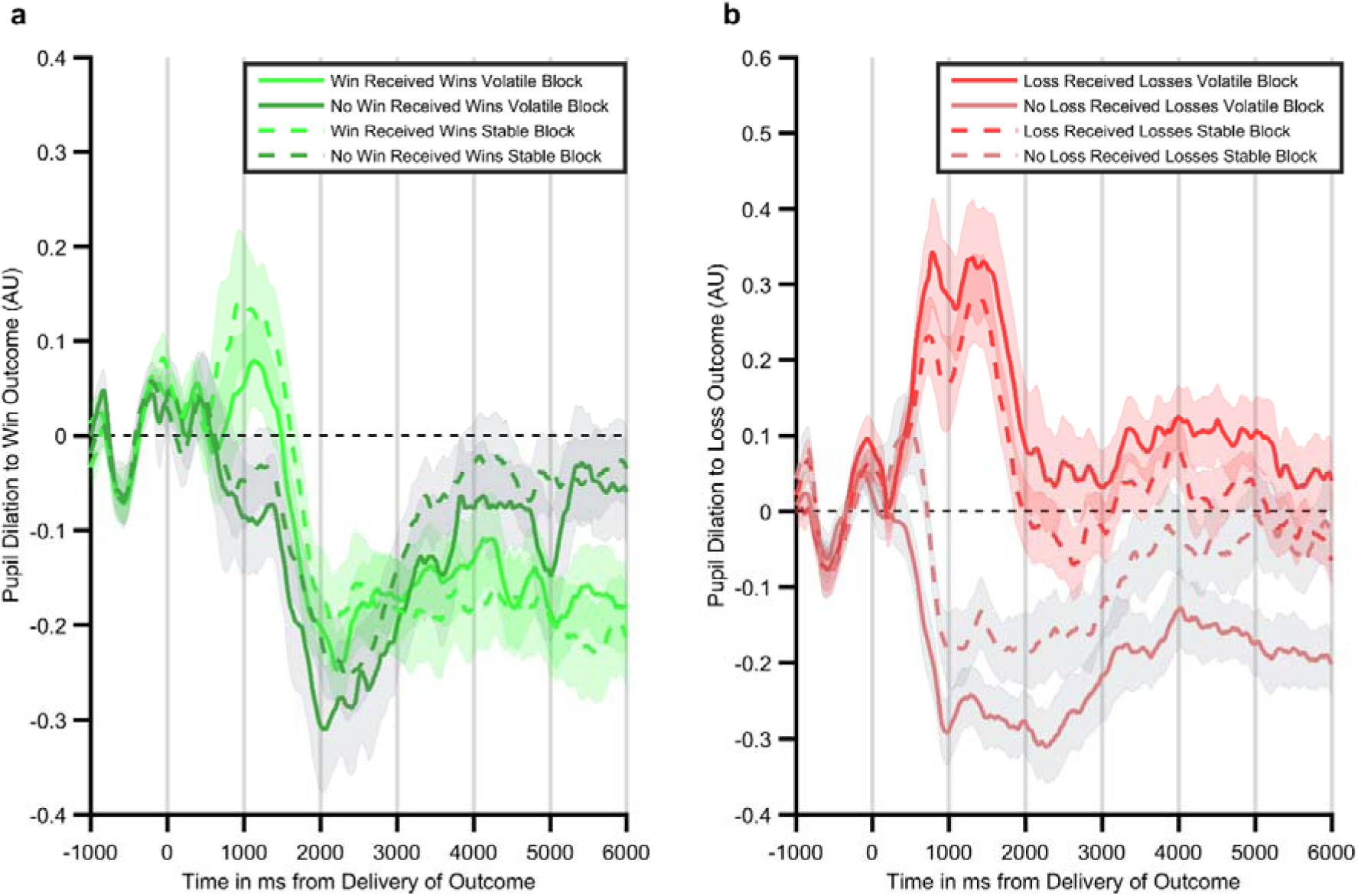
Individual time courses for trials in which wins (panel a) and losses (panel b) are either received or not received. Lines represent the mean and shaded areas the SEM of pupil dilation over the 6 seconds after outcomes are presented.

### Relationship Between Baseline Symptoms of Anxiety and Depression and Task Outcomes

Although participants in the current study were not selected on the basis of their symptoms of depression or anxiety, baseline questionnaires were completed allowing assessment of the relationship between symptoms and task performance. Consistent with previous work (Browning et al., 2015) symptoms of anxiety, measured using the trait-STAI and depression, measured using the QIDS, correlated significantly negatively with differential pupil response to losses (all r<−0.43, all p<0.02). That is, the higher the symptom score, the less pupil dilation differed between the loss informative and loss non-informative blocks. These measures did not correlate with pupil response to wins (all p>0.2). A marginal correlation was found between trait-STAI and the change in learning rate to losses, with participants with higher scores adjusting their learning rate less than those with a lower score (r=−0.34, p=0.07). We did not observe any relationship between either questionnaire measure and change in the win learning rate or between QIDS score and change in loss learning rate (all p>0.2).

